# Concurrent invasions by European starlings (*Sturnus vulgaris*) suggest selection on shared genomic regions even after genetic bottlenecks

**DOI:** 10.1101/2021.05.19.442026

**Authors:** Natalie R. Hofmeister, Katarina Stuart, Wesley C. Warren, Scott J. Werner, Melissa Bateson, Gregory F. Ball, Katherine L. Buchanan, David W. Burt, Adam P.A. Cardilini, Phillip Cassey, Tim De Meyer, Julia George, Simone L. Meddle, Hannah M. Rowland, Craig D.H. Sherman, William Sherwin, Wim Vanden Berghe, Lee Ann Rollins, David F. Clayton

## Abstract

A species’ success during the invasion of new areas hinges on an interplay between demographic processes and the outcome of localized selection. Invasive European Starlings (*Sturnus vulgaris*) established populations in Australia and North America in the 19^th^ century. Here, we compare whole-genome sequences among native and independently introduced European Starling populations from three continents to determine how demographic processes interact with rapid adaptive evolution to generate similar genetic patterns in these recent and replicated invasions. Our results confirm that a post-bottleneck expansion may in fact support local adaptation. We find that specific genomic regions have differentiated even on this short evolutionary timescale, and suggest that selection best explains differentiation in at least two of these regions. This infamous and highly mobile invader adapted to novel selection (e.g., extrinsic factors), perhaps in part due to the demographic boom intrinsic to many invasions.

## Introduction

Some species can establish and spread in a novel environment more successfully than others, and defining what makes a species ‘invasive’ is hotly contested^1-4^. Invasion biologists continue to debate whether an invasion’s success can be better attributed to an intrinsic property of the founding population or to extrinsic conditions experienced by the population. When invasive populations colonize a new environment, they often undergo genetic bottlenecks^5^. However, even populations with limited genetic diversity, including those subject to founder effects, can adapt quickly to novel environments^6-8^. For example, gene surfing during range expansion has led to adaptation in experiments in microbes ^9^ and wild, invasive bank voles ^10^. Simultaneous with demographic shifts, invasive populations may encounter environmental conditions that exert novel selective regimes. Although genetic bottlenecks often limit the genetic variation available to selection, local adaptation can occur even after a genetic bottleneck^6,11-13^.

Recent and replicated invasions are particularly useful for exploring the eco-evolutionary dynamics of population expansion^14-16^, since any divergence after introduction likely reflects a combination of demographic and/or selective forces. One such recent invader is the Common or European Starling (*Sturnus vulgaris*), which was introduced across south-eastern Australia in 1856 and to New York, United States of America in 1890^17^. Both the American and Australian invasions were most likely founded by individuals from the United Kingdom^18,19^, although multiple introductions in Australia might contribute to ongoing gene flow among continents.^20^

The starling’s ecology is well-studied in both invasions, enabling us to explore how environmental factors might impact genetic variation^21^. Native-range starlings thrive in open pastures and urban environments, but starlings’ ecology and life history vary among populations. In range-wide studies of Australian ^22-24^ and North American ^25,26^ invasions, temperature and precipitation influence genetic differentiation, even after controlling for population structure. Migration and dispersal also vary among invasions. In North America, starlings can migrate long distances each season^19,27^, but populations in the Western USA likely disperse and/or migrate shorter distances^28^. In contrast, starlings in Australia exhibit strong population structure and likely migrate short distances in search of food, even though environmental conditions are much more arid than in the native range^22^. Finally, variation in the breeding cycle may also facilitate invasion success as invasive populations tend to lay more than one clutch, whereas the UK population generally lays only one clutch^29,30^.

We use concurrent starling invasions in Australia (AU) and North America (NA) to examine the evolutionary and genetic consequences of invasion. Both starling invasions rapidly expand from small founding populations in the late 19^th^ century^18,19^. Given this natural control, we take advantage of a rare opportunity to compare intrinsic and extrinsic drivers of invasion success. Founder effects and other intrinsic demographic properties of invasions certainly influence establishment ^31,32^, and we predict some divergence from the native, ancestral population (represented by UK samples here) due to genetic drift. We combine demographic models with fine-scale measures of genomic diversity and differentiation to determine whether drift alone can explain observed genetic variation, and we find strong evidence that extrinsic factors such as novel selective pressures generate differentiation in both invasions specific to only a few genomic regions.

## Results

### Differentiation and population structure

We sequenced whole genomes of eight native-range individuals from Northumberland, UK and eight from each of the two invasions (NA: all 8 from New York City, USA; AU: 2 each from four locations in New South Wales, Australia, Table S1). After filtering, we obtained more than 11 million SNPs with minimum 5X coverage (average genome-wide coverage = 18.44, for details on how choice of reference genome impacts variant-calling, see SI Section 1). All patterns identified using a variant-called dataset concur with those based on genotype likelihood (ANGSD); for details on how variant-calling impacts patterns, see SI Section 2. We use a variant set filtered for minor allele frequency, Hardy-Weinberg Equilibrium, and linkage disequilibrium for analyses of population structure, but for more accurate estimates of genetic diversity and differentiation, we report results from a genome scan of variants filtered only for quality and depth. Differentiation of the two invasive populations from the native population is low, which is expected given that these populations split less than two centuries ago. However, differentiation between AU and UK (FST AU vs. UK = 0.016) is almost twice that between NA and UK (F_ST_ NA vs. UK = 0.008). We examined this contrast in genetic differentiation using analyses of population structure.

All three populations are readily distinguished from one another by a principal component analysis of 40,488 unlinked SNPs (minor AF > 0.25): PC1 (6.3% of genetic variation) separates the UK and NA populations from the Australian population (Figure 1A). This evidence complements previous work that showed extensive population structuring in Australia but nearly continuous gene flow across North America, based on reduced-representation genomic data^22,33^. Principal component analysis in the genotype likelihood framework of ANGSD^34^ shows nearly identical results (Figure S6). Furthermore, individuals are reliably assigned to clades based on pairwise genetic distances calculated in ANGSD (Figure 1B).

**Figure 1.**
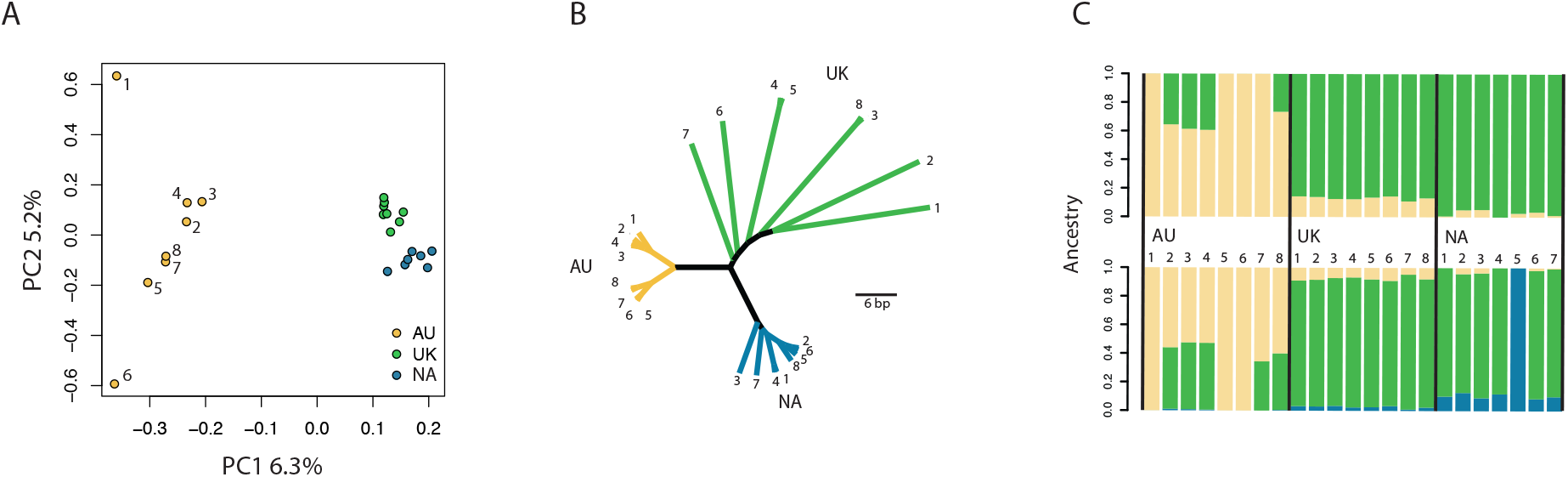
A) Principal components of 40,488 unlinked SNPs explain 6.3% (PC1) and 5.2% (PC2) of genetic variation. B) cladogram of genetic distances among samples based on genotype likelihoods of 16,151,007 sites. C) ADMIXTURE analyses showing K=2 (top) and K=3 (bottom row).

We note that the tight clustering of UK individuals in Figure 1A contrasts with the large distances between these same individuals in Figure 1B. In the genotype likelihood dataset, genetic distances between native UK individuals are much greater than distances among individuals within each invasive population (Figure 1B). Because these two datasets differ in variant-calling and filtering strategies, the genetic distance among UK individuals in Figure 1B may reflect rare alleles that were filtered out of the variant set in Figure 1A. However, PCs 3 and 4 in the PCA of the variant-called dataset do indicate additional structure within the UK population (Figure S7).

Admixture analyses revealed statistical support for a two-population model, which distinguished AU from NA+UK and was only slightly weaker (cross-validation error = 0.88) than the best-supported model of K=1 (cross-validation error = 0.73, Figure 1C, SI Section 3). Regardless of the number of populations hypothesized in models of admixture, the NA invasion consistently shares a higher proportion of its ancestry with the UK population. Individuals from the Australian population are distinguished from the other two populations in all tested values of K. Considered in concert, these tests of population structure show that Australian and North American populations differ in the amount of divergence from the native UK population. Founder effects likely contribute to the observed population structure, and below we describe explicit models of demographic processes.

### Invasive and native populations experienced bottlenecks and subsequent expansion

To address the impact of demographic processes in generating the observed patterns we used the site-frequency spectrum built from genotype likelihoods to construct models of changes in the effective population size over time. We examined the demographic history of all three populations using fastsimcoal2 ^35^. The demographic model shows that both invasions experienced a bottleneck upon introduction (Figure 2), which is exactly what we expect given the small number of founding individuals in both AU and NA. Bottlenecks often lead to inbreeding within a population, and we find that inbreeding is negligible but slightly higher in the Australian population than in the NA population (Table S2). In addition, relatedness among individuals is quite low, where zero indicates no shared alleles among individuals (maximum AJK statistic = 0.06).

**Figure 2.**
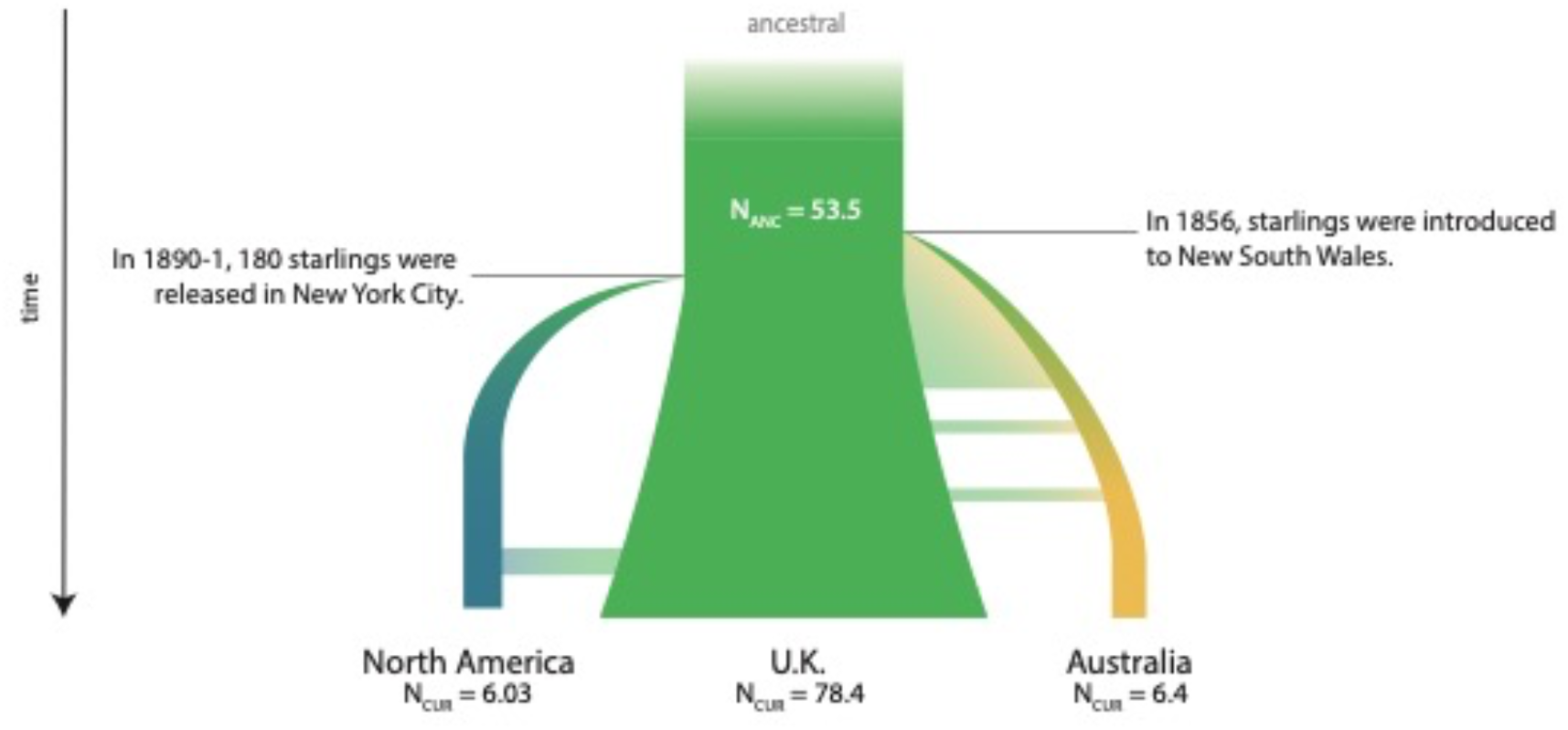
Demographic model of effective population size based on the site-frequency spectrum. Schematic approximates population growth based on model output from *fastsimcoal2*.

Each invasion appears to have recovered quickly by expanding in effective population size, and at present, our data suggest similar effective population sizes in both the AU and NA invasions. Much of the ancestral variation appears to be shared among invasions, given that the genome-wide average F_ST_ between AU and NA is 0.04, although we note that genetic differentiation among invasions confirms the expectation that different variants will make it through each genetic bottleneck. The results presented here concur with range-wide sampling that indicates genetic bottlenecks followed by rapid expansion with little evidence of inbreeding in both Australia^22^ and North America^33^.

Population bottlenecks often lead to a loss of genetic diversity when genetic drift drives to fixation alleles once maintained at a moderate frequency. Since both the AU and NA invasions experienced a genetic bottleneck, we expect that nucleotide diversity (π) within each population should be higher in the older and larger native range population than in each invasive range. However, we find that 70% of 50-kb windows show greater nucleotide diversity in AU (compared to UK diversity), and 74% show greater diversity in NA. The filters applied should not lead to disproportional allelic dropout of rare alleles, because we did not filter for minor allele frequency prior to this scan. Higher invasive diversity may be a sampling artifact: rare variants in the native range may have ‘surfed’ to a higher frequency in the invasions, and our sampling of the native range (in Northumberland, UK) likely represents only a small portion of genetic variants that could have been introduced in AU and NA. We also note that the UK individuals sampled here do not capture range-wide diversity in the native population, and therefore we expect that actual nucleotide diversity in the UK population is higher than what we have sampled here. Additional investigation of native range starlings will be needed to determine whether lower diversity in the UK is recovered with broader geographic sampling.

Our demographic models show that both the AU and NA invasions rapidly expanded in population size after the initial bottleneck, which predicts that many loci would be lost upon establishment of the invasive populations even as variants in other loci increase in frequency. If gene surfing facilitated by rapid population expansion explained genome-wide differentiation, then we would expect to find that regions where invasive diversity is higher than diversity in the native range are distributed across many chromosomes. However, shifts in diversity and differentiation that occur only in a few narrow regions of the genome would suggest evolutionary dynamics specific to that region. If regions that have differentiated (e.g., high F_ST_) from the native range also show higher diversity in the invasions, this may suggest either a relaxation of purifying selection or an increase in diversifying selection at specific regions of the genome. We note here that the recombination landscape may also contribute to these patterns, and discuss this factor in the following section.

### Differentiation of a few genomic regions reveal consequences of invasion

Starling populations colonized their invaded ranges less than two centuries ago, and the age of these populations makes it somewhat surprising to find loci specific to the derived populations with fixation indices as high as 0.57 in a single 10-kb window (Figure 3). However, no putative outlier windows approach fixation in any comparison across all individuals in a population. We consider only the top 0.1% of 50-kb windows to be F_ST_ outliers (AU vs. UK: F_ST_ > 0.30; NA vs. UK: F_ST_ > 0.22); subsequent references to outliers indicate these windows. Outliers include Chromosomes 1, 2 and 6, and to a lesser extent, 1A, 4, and 4A. Examining localized shifts in diversity and differentiation at these regions can help to resolve the relative contributions of demography and selection.

**Figure 3.**
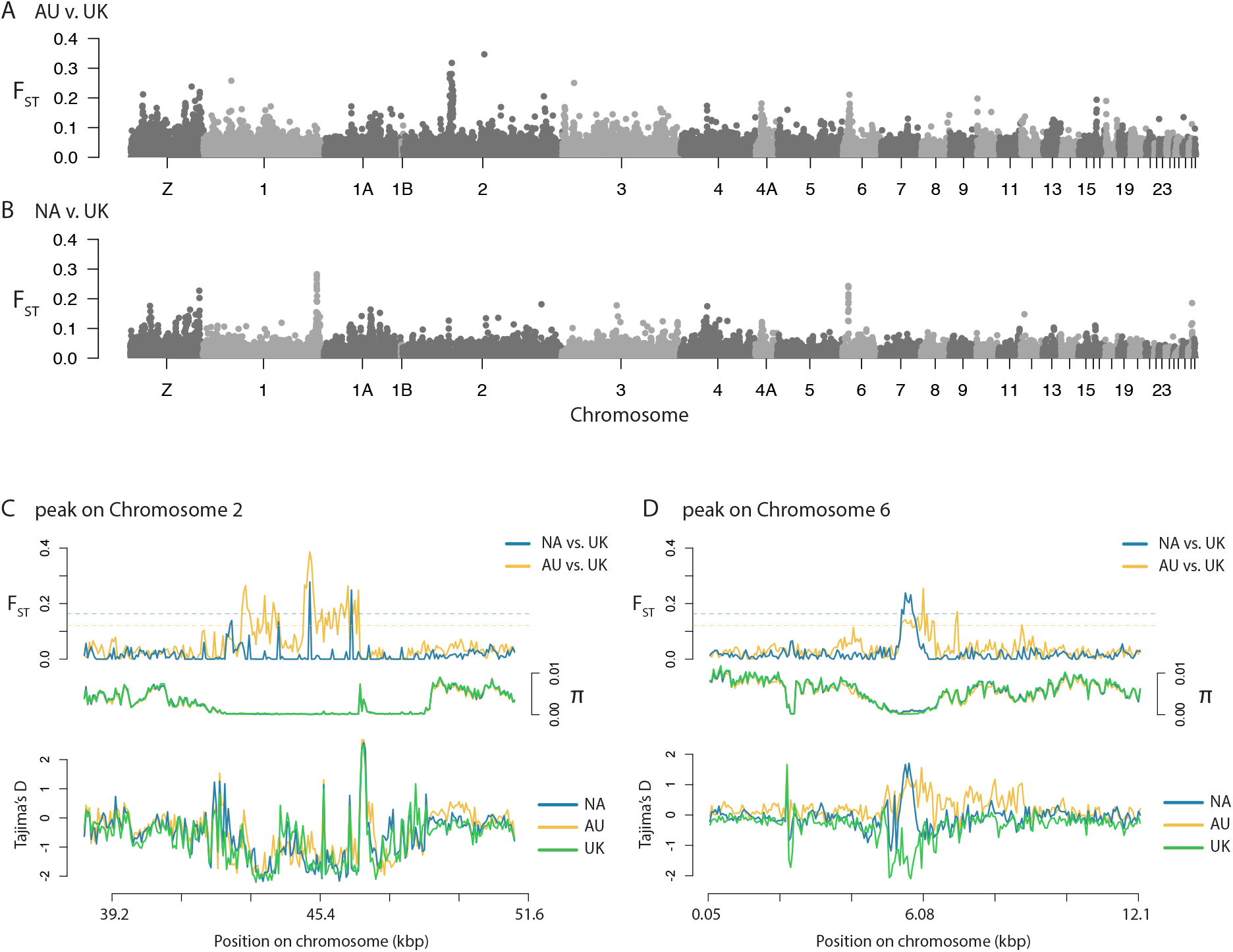
A-B) Manhattan plots show 50-kb windowed F_ST_ between AU & UK (A) and NA & UK (B) populations. C-D) 50-kb windowed F_ST_ (top row), π within each population (middle), and Tajima’s *D* within each population (bottom row), centered on the elevated F_ST_ regions of Chromosome 2 (C) and Chromosome 6 (D). Color represents each population, except in F_ST_ plot where yellow indicates F_ST_ AU vs. UK and blue indicates F_ST_ NA vs UK.

Genome scans like this one tend to search for genomic islands of divergence, where recombination can explain divergence among populations^36^. High levels of differentiation paired with a reduction in diversity may stem from suppressed recombination (e.g., proximity to the centromere or a structural rearrangement), or from alleles approaching fixation or loss due to drift or selection (e.g., a selective sweep). We find that most putative outlier regions are distant from the centromere location (predicted via homology with the zebra finch (*Taeniopygia guttata*) genome, Supplementary Information). Nevertheless, genomic architecture—including linkage disequilibrium independent of the centromere—plays a role in how differentiation among populations is generated. Below, we discuss how demography and selection drive divergence in invasions, considering the context of the starling’s genomic architecture.

### Some F_ST_ outlier regions show clear signals of selection in one or both invasions

If directional selection were driving differentiation between an invading population and its native ancestral population, we would expect to see a decline in nucleotide diversity specific to the invading population. But, as described above, local reductions in π could also result from population bottlenecks experienced during founder events; to clarify the impact of selection, we look Tajima’s *D*. We infer directional selection where most variants are either very rare or very common (e.g., negative Tajima’s *D*). We examine these predictions by looking at two F_ST_ peaks on Chromosome 1A; in this region, we find very low nucleotide diversity and negative Tajima’s *D* in both AU and NA, which is exactly the signature we would expect under purifying selection (Figure S12). Around that region, Tajima’s *D* in the UK population varies between 0 and 1. We note that genetic distance (d_xy_) could also clarify mechanisms, but as d_xy_ tracks F_ST_ perfectly, we argue that linked selection offers the clearest explanation. This pattern is echoed on Chromosomes 4 and 4A, but the remaining outlier regions show more complicated signals (Figures S13-14).

Where F_ST_ is highest on Chromosome 2, we find strong evidence of both purifying and balancing selection in all three populations (Figure 3C). We find that nucleotide diversity is very low within every population, and immediately after the block of elevated F_ST_, we see a sharp increase in nucleotide diversity in all three populations (Figure 3C). In addition, Tajima’s *D* tracks these changes in diversity: where F_ST_ and π are low, Tajima’s *D* is also negative in all three populations—indicating an excess of low-frequency variants and perhaps purifying selection—but this statistic climbs to high positive values immediately before and after the block of elevated F_ST_, indicating balancing selection. The concordance of Tajima’s *D* in all three populations suggests a release of some kind, whether it is a relaxation of purifying selection or a recombination breakpoint. Even though the centromere is predicted to be 30-Mb downstream of this region, these signatures are consistent with linkage disequilibrium in this 4-Mb region: eukaryotes generally show suppression of recombination near the centromere, leading to a build-up of linkage disequilibrium if this suppression extends for 30-Mb. It is possible that a structural variant in the founding population could generate this pattern. However, we note that F_ST_ among AU and the other populations in fact declines dramatically (to around 0.1) in the middle of the 4Mb region, and in the same location, F_ST_ between NA and UK increases slightly. If this genomic region differentiated as a large linkage block, we would not expect such a decline in F_ST_ and a weakening of selective pressure (as evidenced by the increase in Tajima’s *D*). For these reasons, we suggest that the peak on Chromosome 2 indicates both purifying and balancing selection in the AU invasion.

### One region on Chromosome 6 reveals how population expansion could interact with selection

Both invasions have differentiated from the native range in a 4-Mb region of Chromosome 6 (Figure 3). As a preliminary check, we note that the large distance between this F_ST_ peak and the centromere suggests that low recombination is unlikely to explain differentiation. We suggest that the clearest explanation of this peak invokes selection on previously rare variants, based on three lines of evidence.

First, we suggest that rapid population expansion allowed previously rare variants to surf to a higher allele frequency in the invasions. In this 4-Mb region, invasive diversity (π_AU_ and π_NA_) are each more than three times the native diversity. This shift in within-population diversity is not random; in fact, when we examine invasive nucleotide diversity directly under the F_ST_ peak, we find only three other windows ∼4.2-Mb upstream of this peak show invasive diversity that is notably higher than native diversity. This evidence supports the hypothesis that upon establishment, starlings experienced either (1) balancing selection (strong positive Tajima’s *D*) in both invasions due to novel selective pressures or (2) a release of purifying selection that led to an accumulation of variants and thus higher invasive diversity—but only in this specific region. These patterns could be driven by a small number of individuals, or they could indicate a population-wide shift, which leads us to our next point.

Second, in this region, we find that these higher-diversity alleles in both the NA and AU populations have increased in frequency relative to the native range. In the same region, we find strong positive values of Tajima’s *D* in the invasions—indicating a moderate-to-high frequency of the alternative allele—and negative Tajima’s *D* in the UK population at this peak, since these signatures suggest that previously rare variants have increased in frequency in the invasions only. Alternatively—or simultaneously— purifying selection may have driven these same variants to a lower frequency in the UK population. The most parsimonious explanation of these shifts in diversity is a single event in the UK population, but we note that this shift is specific to only a small region of the genome.

Third, and most importantly, these patterns are found in this region of the genome only, and it is notable that this shift in diversity co-occurs with one of the highest F_ST_ peaks. We would expect to see similarly high invasive diversity under other F_ST_ peaks if population expansion alone could explain these patterns. However, nowhere else in the genome do we find such high invasive diversity where native diversity is low. We suggest a selective explanation given that a genetic bottleneck is not likely to produce this pattern. Taken together, these results provide evidence that rapid expansion of these starling invasions may have facilitated selection to drive previously rare variants higher in frequency, independently in both NA and AU populations.

### Genes under putative selection may aid in invasion success

The region under putative selection on Chromosome 6 overlaps with the coding regions of four genes with dramatically different functions (*JMJD1C, RTKN2, NRBF2*, and *ARID5B*), and we suggest that selection on one of these genes might explain the differentiation in the region, with the other genes remaining in linkage disequilibrium with the possible candidate. Among these candidates, we can speculate that *ARID5B* has the most intuitive link to hypothesized selective pressures: this protein is required for adipogenesis and involved in smooth muscle differentiation. The first exon of this gene lies directly under the F_ST_ peak between AU and UK starlings, and muscle growth and fat storage may have been key to dispersal ability. The three other genes that overlap this window are involved in the DNA-damage response (*JMJD1C*), lymphopoiesis (*RTKN2*), and regulating autophagy (*NRBF2*). For details on GO term enrichment in these outlier regions, see SI Section 6. Regardless of the mechanism driving these loci toward an intermediate frequency, it remains possible that variation at one or more of these loci influenced invasion success by maintaining heterozygosity.

### Conclusion

An open question in invasion biology is whether an invasive population’s success is better attributed to intrinsic properties of the invasive species or to extrinsic factors specific to the novel environment. These results demonstrate how demographic shifts during and immediately after any establishment may support a species’ success and even lead to local adaptation to environmental conditions. The Australian and North American starling invasions colonized each continent around the same time, and experienced similar contractions in population size that led to classical founder events upon establishment. The shared decline in genetic diversity represents a shared intrinsic determinant of invasion success. However, differences in propagule pressure and dispersal likely influence the evolutionary trajectory of each population ^37,38^.

Propagule pressure (also termed introduction effort) is a composite measure of the number of individuals released, and we note that founding population sizes may have varied slightly among AU and NA introductions. Even without dramatic variation in founding population sizes, dispersal itself shapes genetic diversity and thus adaptive potential, as shown in a recent study of invasive plants ^39^. Starlings’ dispersal and migration varies among populations: starlings in Australia tend to migrate less frequently and across shorter distances, but starlings migrate and/or disperse hundreds of kilometers in North America ^19,40^ and South Africa^41^.

It is remarkable that despite this contrast in life history strategies and the stochastic nature of evolution during range expansion, we find F_ST_ peaks shared among invasions at only a few regions of the starling genome. Although these F_ST_ peaks could arise via drift, footprints of other population genetic metrics are consistent with selection. We note that mutations in specific chromosomal regions could also be accelerated by extrinsic environmental properties (climate, food availability, and more) through epigenetic CpG DNA hypermethylation events, which are known to increase frequency of genetic mutations by spontaneous deamination CG>TG transition ^42-44^. For example, such epigenetic shifts supported the invasion of another avian species (the house sparrow) into Australia^45^. Regardless of genetic mechanism, we suggest that differentiation in these genetic regions is simultaneously shaped by intrinsic and extrinsic drivers of invasion success. We find it notable that some differentiated regions (in particular, Chromosome 6) are shared among invasions despite differences in the selective environment as well as stochastic processes that shape the starling’s evolution on each continent. The European starling invasions compared here suggest that rapid population growth may support local adaptation.

## Methods

### Whole-genome re-sequencing

Libraries for each individual starling were constructed using a TruSeq DNA PCR-free High Throughput Library Prep Kit (Illumina, San Diego, CA). All individuals passed the initial quality check with FASTQC (Babraham Bioinformatics, Cambridge, UK).

Adapters were removed using AdapterRemoval^46^ and reads mapped to the reference *S. vulgaris* vNA genome (GCF_001447265.1)^47^ genome using BOWTIE2^48^ and checked for mapping quality using qualimap^49^. Sequencing quality was relatively high: 96.4% of reads mapped to the *S. vulgaris* genome with a coverage of 18.4X and a mapping quality of 26.9 (Table S1). Reads were also mapped to a pseudo-chromosome-level *S. vulgaris* vNA genome, where scaffolds were assigned to chromosomes based on orthology to the zebra finch reference genome (GCF_000151805.1)^50^. Assuming orthology, we were able to predict centromere positions based on the known genomic architecture of the zebra finch^51^, but this study does not directly define centromere position.

We called variants using GATK’s HaplotypeCaller in GVCF mode and flagged low-quality variants using GATK Best Practices (QD<2, FS>60, MQ<40, and SOR>3, accessed March 21, 2018 ^52^). We filtered sites for missing data, depth, and quality using vcftools (parameters: --max-missing 0.8 --min-meanDP 2 --max-meanDP 50 --remove-filtered-all), which removed 4.1 million sites from the original SNP set and left a total of 23.4 million sites for downstream analyses. Starting from the mapped reads used in the GATK pipeline, we also called SNPs based on a minimum p-value of the correct genotype probability at each site using ANGSD^34,53^. Filtering for SNP p-value (0.0001), depth (between 60 and 400 sequences), and mapping quality (>20) left 16,151,007 sites. All scripts used in read processing and filtering are available on GitHub: https://github.com/nathofme/global-RESEQ.

### Population structure

Population structure analyses used a dataset of biallelic SNPs in Hardy-Weinberg Equilibrium, where minor allele count (MAC) is > 2 and SNPs were pruned for LD by removing all sites with an r^2^ > 0.6 within 1kb sliding windows. This filtering left 868,685 sites. Since some individuals showed much lower coverage (minimum 5.58X), all tests of population structure were run with both variant-called (GATK) and genotype probability (ANGSD) datasets. Scripts for variant-called analyses are stored at https://github.com/nathofme/global-RESEQ/blob/master/filter-scan.sh, and scripts for probability-based analyses at https://github.com/nathofme/global-RESEQ/blob/master/angsd.sh.

We estimated variance among and between individuals using a principal components analysis in SNPRELATE^54^ (GATK) and a covariance matrix built in ngsTools^55^ (ANGSD). We used ADMIXTURE ^56^ (GATK) and NGSADMIX^57^ (ANGSD) to examine shared ancestry among individuals, and we also measured pairwise genetic distances using ngsDist for the ANGSD dataset only.

### Demographic inference

We used FASTSIMCOAL2^35^ to explicitly test for genetic bottlenecks in each population. FASTSIMCOAL2 takes a site-frequency spectrum (SFS) as input, and we used the SFS estimated from ANGSD given that likelihood-based estimates are more robust to sequencing error^58^. Demographic models in *fastsimcoal2* used priors on the time (TBOT = 10 to 300) and size of the bottleneck (NBOT = 10 to 1000). The command line arguments were as follows: -M -n 1000000 -L 50 -q -k 100000. Each run began with a randomly generated seed (-r), and the -k flag simply writes polymorphic sites to a temporary file to cope with the high memory usage of this analysis. Scripts for demographic analyses can be found at https://github.com/nathofme/global-RESEQ/blob/master/demography.sh.

To verify this demographic model, we also estimate inbreeding coefficients (F-statistics) using the –het command in VCFTOOLS, and calculate relatedness among each pair of individuals using the –relatedness method of Yang et al. (2010) in VCFTOOLS.

### Sliding window scans

For scans of genetic divergence and diversity, we used a variant dataset filtered only for depth and quality: we kept variants that had less than 20% missing data across all sites, but did not apply minor allele frequency (MAF) filters, filter for HWE, linkage, or any other factors. Given that rare alleles likely provide the strongest evolutionary signals in this system, we did not want to filter out any alleles that might have been rare in one population (e.g., the native UK population) and increased in frequency in another population (e.g., the AU or USA invasions). Nevertheless, we test for sensitivity to this filtering choice in SI Section 2B.

We calculated F_ST_ and nucleotide diversity (π) using overlapping 50-kb sliding windows with a step size of 10kb using VCFTOOLS ^59^. We calculate nucleotide diversity separately for each population. We then calculate d_xy_ using a Python script by Simon Martin (accessed at https://github.com/simonhmartin/genomics_general/blob/master/popgenWindows.py on February 12, 2020). This analysis includes all confident variant calls (parameters: -- output-mode EMIT_ALL_CONFIDENT_SITES in GATK’s HaplotypeCaller); only with this modification do we recover levels of d_xy_ similar to other systems. We also measured F_ST_ and π in overlapping 10-kb windows to localize elevated F_ST_ to an even smaller region of the genome. To visualize relationships between diversity metrics, we plotted the mean values of each metric in a 50-kb window. All scripts for plotting are stored on GitHub: https://github.com/nathofme/global-RESEQ/.

### Identifying candidate genes under selection

In contrast to identifying single genes, network analyses of gene ontology (GO) terms can provide a more holistic and objective method of identifying shared functions. Network analyses can also provide more statistical power, correcting for the usual problem of multiple testing. To identify functions of candidate regions, we quantified the uniqueness and dispensability of each GO term using a method that quantifies semantic similarity^60^. This analysis by default emphasizes GO terms that are rare in the list of candidates provided; because we are interested in functions that are common across outlier regions, we manually curate category labels to choose GO terms that are less unique and more dispensable as representative titles.

## Acknowledgments

SLM acknowledges Roslin Institute Strategic Grant funding from the UK Biotechnology and Biological Sciences Research Council (BB/P013759/1). HR acknowledges that sequencing costs were supported by a fellowship from The Winston Churchill Memorial Trust, and a Phyllis and Eileen Gibbs Fellowship from Newnham College, Cambridge. LAR was supported by a Scientia Fellowship from UNSW. This project was also supported by discussions at Bird Sense meetings funded by the Royal Society of London, the Association for the Study of Animal Behaviour, and the Zoological Society of London. NRH acknowledges bioinformatic support from members of the Fuller Evolutionary Biology Program at the Cornell Lab of Ornithology, including Irby Lovette, Jennifer Walsh, and Leonardo Campagna.

## Author Contributions

Project conception: all authors

Field work: Melissa Bateson, Lee Ann Rollins, Adam Cardilini, Scott J Werner

Lab work: Wes Warren and the McDonnell Group at WUSTL.

Data analysis and manuscript writing: Natalie R Hofmeister, Katarina Stuart

Manuscript editing: all authors

## Notes

### Competing Interest Statement

The authors have declared no competing interest.

