## Supplementary Material for "Concurrent invasions by European starlings (*Sturnus vulgaris*) suggest selection on shared genomic regions even after genetic bottlenecks"

### 1. Three different genome assemblies produce similar results

We first compared how different genome assembly versions may affect variant calling and further downstream analysis. The raw data from a global whole genome starling comparison project was realigned to each of the three genome assemblies. A scaffolded version of *S. vulgaris* vNA (GCF_001447265.1, scaffolded using Zebra Finch genome GCF_008822105.2 and Satsuma2), *S. vulgaris* vAU1.0^1^, and lastly a non-scaffolded version of *S. vulgaris* vAU that involved an initial supernova (v2.1.1) ^2^ assembly and polishing using Pilon (v1.23)^3^ of the 10x linked-read data in Stuart and Edwards *et al.* 2021. The variant calling pipeline was similar to that used in the main manuscript, with some program version differences. Briefly, eight *S. vulgaris* individuals from each of the United Kingdom (UK; native range), North America (USA; invasive range), and Australia (AU; invasive range) were sequenced using short read Illumina sequencing. The raw reads were processed using samtools (v1.9)^4^ and bedtools (v2.27.1)^5^, and adapters were removed using adapterremoval (v2.2.2) ^6^. The processed reads were aligned to each of the three genome assemblies (Table 1) using bowtie2 (v 2.3.5.1) ^7^, and indexed using picard (v2.18.26)^8^. gatk’s (v4.1.0.0) ^9^ HaplotypeCaller (GVCF mode) was used to call haplotypes for each sample, which were processed with CombineGVCFs and finally passed into GenotypeGVCFs to produce the initial VCF file. The VCF file was put through an initial filter step in gatk (QD<2.0, FS>60.0, MQ<40.0, SOR>3.0), and a secondary filter step (max missing count=4, min mean DP=2, max mean DP=50, min alleles=2, max alleles=2). VCFtools (v0.1.16) ^10^ was then used to create three sets of filtered VCF files for each genome. A SNP data set was created by filtering for minor allele frequency (MAF) of 0.1, and one data set was created at MAF 0.05. The MAF 0.05 dataset was then further filtered; SNPs were pruned in bcftools (samtools v1.9) for linkage by removing sites with an r^2^ > 0.6 within 1000 bp site windows ^11^. This last data set was used for population genetics analysis.

The three variant data files were assessed using samtools (v1.9) *bcftools* *stats* function. The final data set (MAF 0.05 and LD filtering) was parsed through snprelate (v1.22) ^12^, and displayed in a PCA to illustrate individual clustering and population relatedness. admixture (v1.2) ^13^ analysis was used to explore population relatedness and admixture.

Some minor differences in SNP counts and the levels of SNP missingness per individual was found between the non-scaffolded *S. vulgaris* vAU genome version and the other two scaffolded genome versions (Fig. 1, Fig. 2). This difference is likely due to smaller scaffold sizes affecting the mapping of the short whole genome sequencing reads in the non-scaffolded *S. vulgaris* vAU genome version. However, these discrepancies were minor, and no biologically meaningful differences were found in the population differentiation analysis using the three different genome versions (Table 1, Fig. 3, Fig. 4). These results indicate that neither the population from which the reference individual has been sourced, nor the level of genomic scaffolding (continuity of the genome), has an effect the SNP based population analysis conducted above. Further, non-scaffolded *S. vulgaris* vAU did not undergo chromosomal alignment to the chromosomes of the *T. guttata*, while the other two genome versions used here did undergo this scaffolding process. This indicated that the synteny alignment conducted had no biological impact on the population differentiation analysis conducted, and all analyses in the main text and the rest of this supplement use the vNA version.

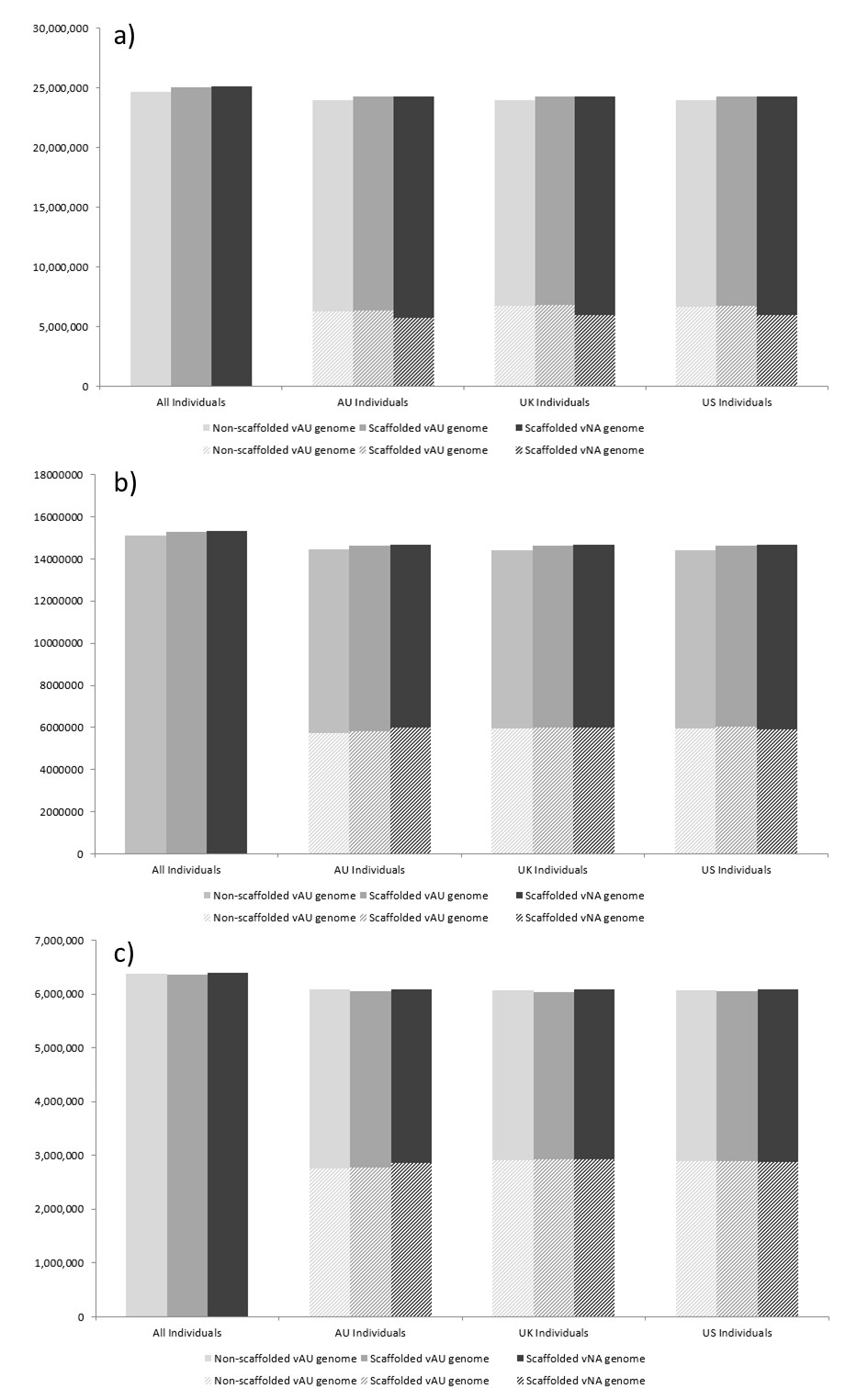

**Figure S1.** Number of alleles recovered for each of the three *S. vulgaris* genome versions. a) MAF of 1%, b) MAF 5%, c) MAF 5% and linkage filtered at r2<0.6 in 1000 bp sliding windows. Total alleles (solid) contains both reference and non-reference alleles (nRefHom + nNonRefHom + nHets), while non-reference SNP (stripped) is a nNonRefHom and nHets (nNonRefHom + nHets). Counts are obtained from bcftools, SNP counts for AU, UK and NA individuals were obtained by calculating the values per individual and averaging them across the population (n=8).

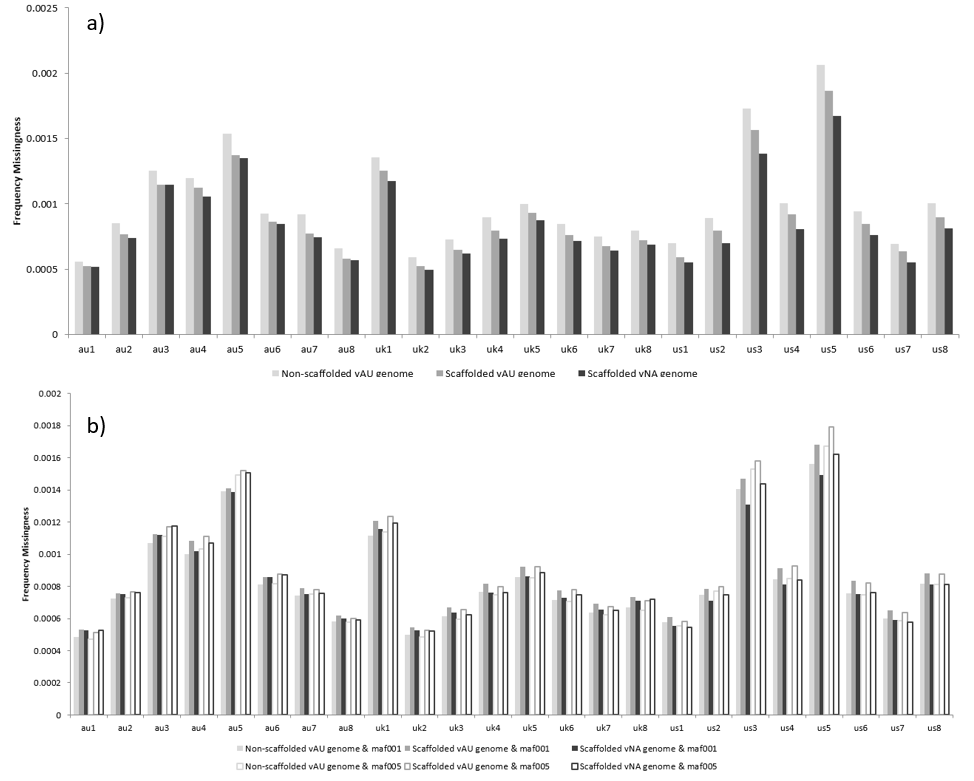

**Figure S2.** Missingness per individual for each genome assembly. a) MAF 5% and linkage filtered at r2<0.6 in 1000 bp sliding windows, and b) MAF 1% (dark) and MAF 5% (light). Individual missingness calculated using vcftools.

**Table S1.** Differences in genome-wide F_ST_ among genome assemblies. F_ST_ calculated between pairwise population comparisons (n=8) for MAF = 0.05 and r^2^<0.6.

| **Non-scaffolded vAU genome** | | | |
| --- | --- | --- | --- |
|  | **AU** | **UK** | **NA** |
| **AU** | - | 0.0431668 | 0.045913 |
| **UK** | - | - | 0.0374013 |
| **Scaffolded vAU genome** | | | |
|  | **AU** | **UK** | **NA** |
| **AU** | - | 0.0430631 | 0.045771 |
| **UK** | - | - | 0.0373351 |
| **Scaffolded vNA genome** | | | |
|  | **AU** | **UK** | **NA** |
| **AU** | - | 0.043003 | 0.045816 |
| **UK** | - | - | 0.037413 |

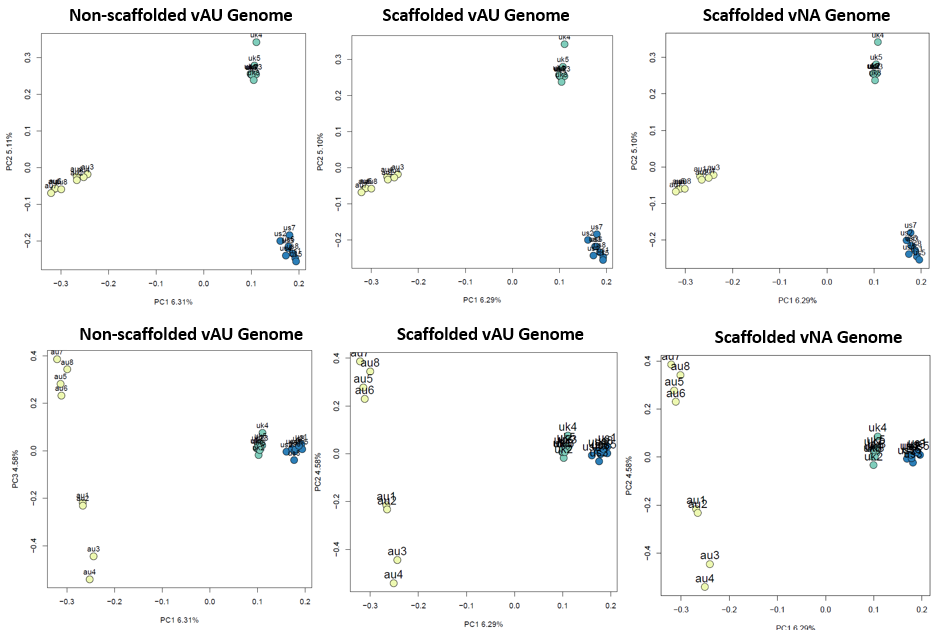

**Figure S3.** PCA comparisons among genome assemblies. The variant data are filtered using MAF 5% and linkage threshold of r2<0.6 in 1000 bp sliding windows. Produced using snprelate.

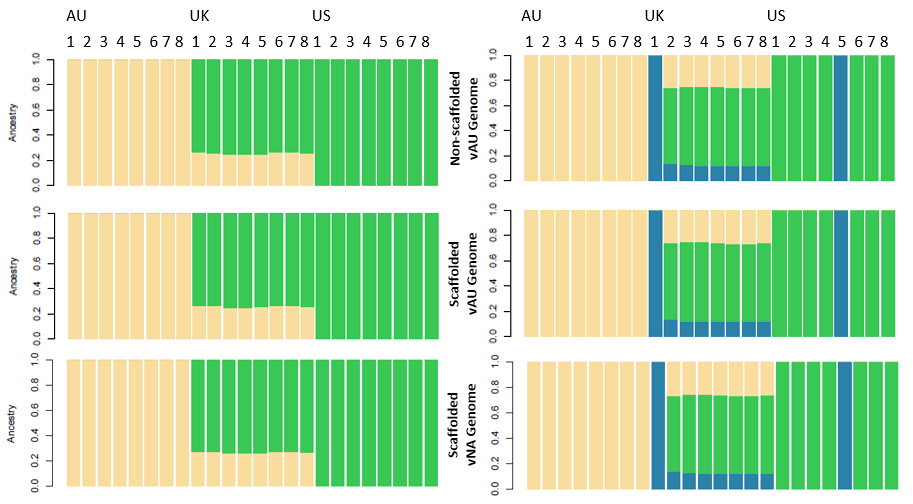

**Figure S4.** Admixture comparisons among the three *S. vulgaris* genome versions. K=2 is shown in the left column and K=3 in the right column. These data are filtered using MAF 5% and linkage filtered at r2<0.6 in 1000 bp sliding windows.

### 2. Variant-calling method does not change results dramatically

#### A. Sequencing data quality

Recent advances in genomic methods urge caution when calling and filtering variants, and we note that the methods we use here represent only one of many possible strategies. Representing variants in the form of likelihoods captures the uncertainty inherent in sequencing data, and this approach is essential when working with low-coverage or low-quality data. However, when sequencing depth and quality is relatively high, we can be more confident that the called variant is the true variant at that site. In these data, sequencing depth was relatively high even before filtering (Figure S5), but we describe our sensitivity analyses among variant-calling approaches in Section 2B below.

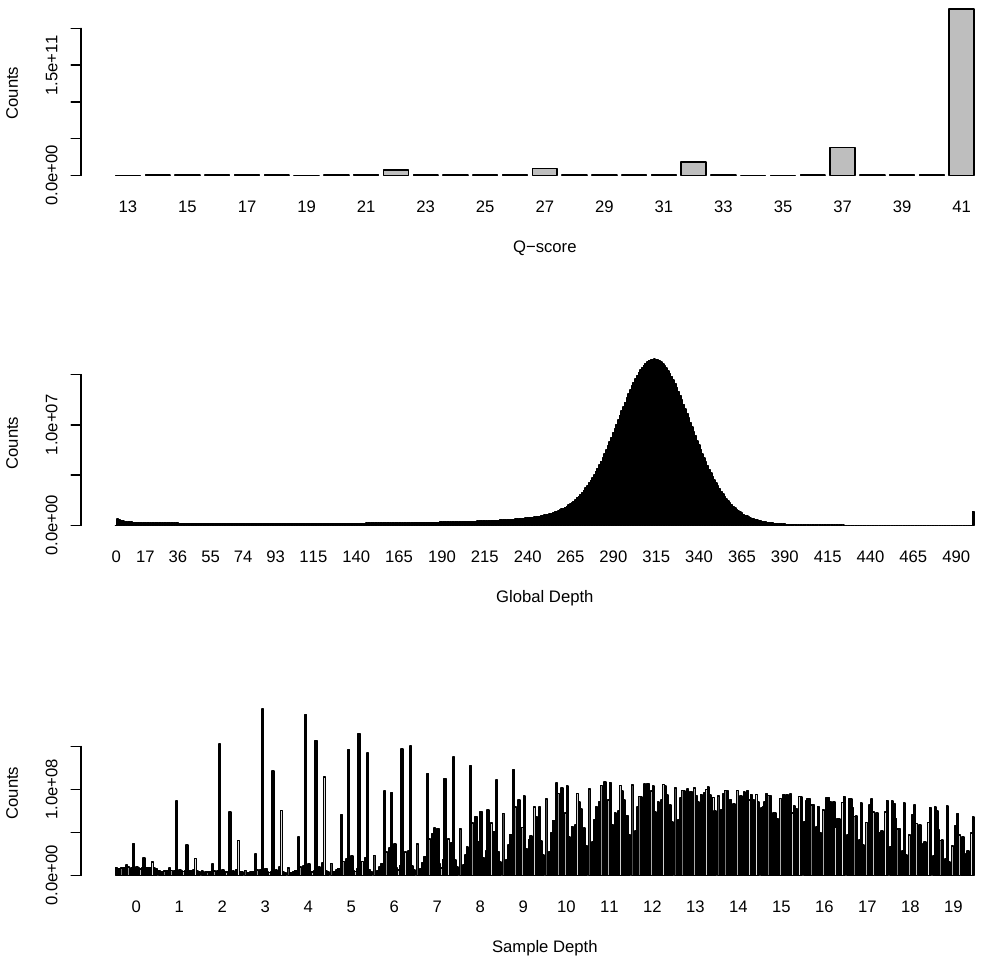

**Figure S5.** Quality of sequencing data as quantified in ANGSD. Top row indicates base quality score (Q-score), middle row shows read depth across all individuals (global depth), and bottom row shows sequencing depth for each individual (sample depth.)

#### B. Impact on diversity metrics

First, we note that genome-wide patterns are sensitive to whether or not invariant sites are included in genome scans: although the generalized results presented here are consistent regardless of which sites are included, F_ST_ can change dramatically when invariant sites are included or not. By way of example, here is a qualitative description of how patterns of differentiation on Chromosome 2 change: in a sliding scan of 50-kb windows, F_ST_ peaks in US vs. UK show up with variant sites only but not when we include invariant sites.

In addition, minor allele frequency filters

#### C. Impact on population structure

We first checked whether population structure results were sensitive to variant-calling method. We find that a principal components analysis using genotype likelihoods yields results consistent with the variant-called set in Figure 1 of the main text.

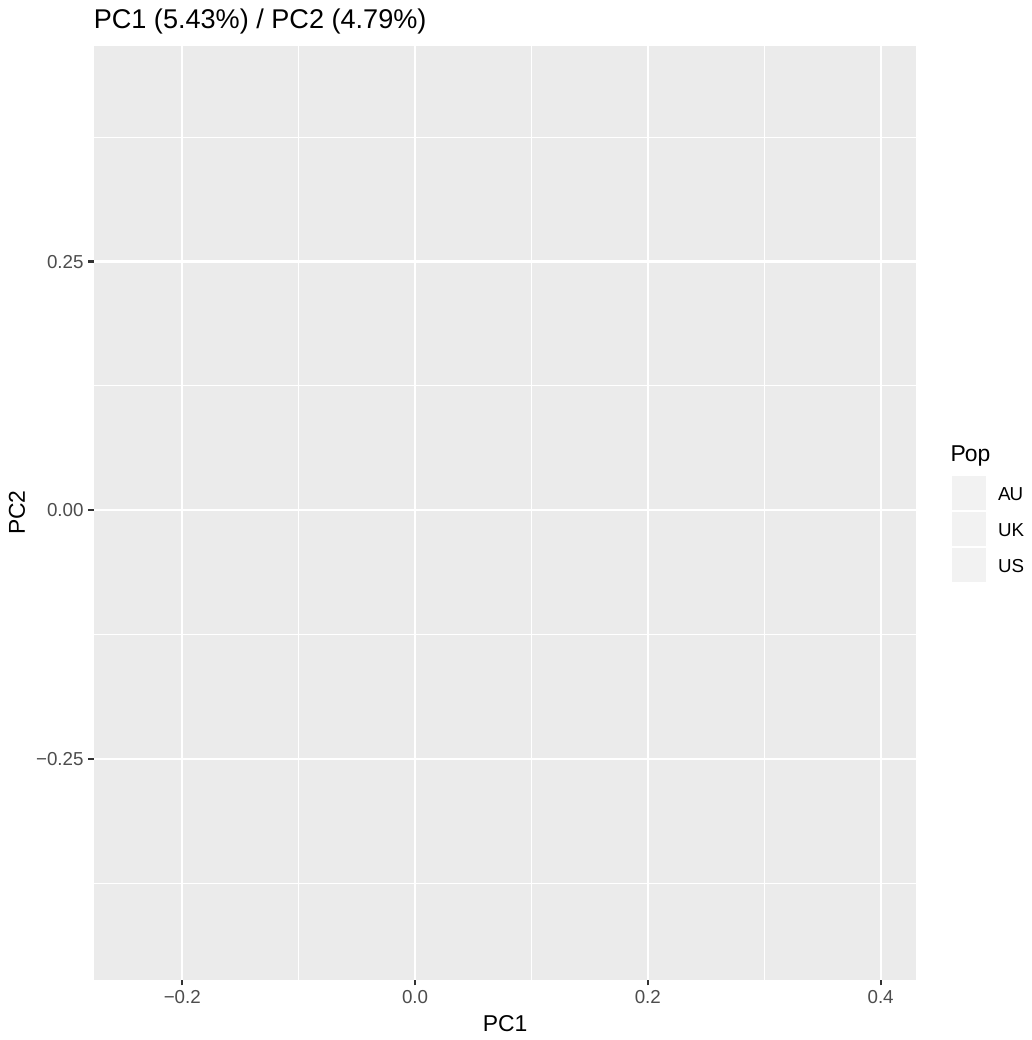

##### **Figure S6.** Principal components analysis of the same dataset using genotype likelihoods, to compare to called variants in the main text.

#### D. Impact on demographic model

Demographic models rely on subtle differences in allele frequencies to infer bottlenecks or other demographic shifts, but these differences can easily reflect sequencing errors. In particular, singleton frequency is highly dependent on sequencing quality. For this reason, any demographic methods presented in this manuscript use a site-frequency spectrum built within the ANGSD framework to avoid possible biases due to sequencing errors.

### 3. Additional tests of population structure

PC2 distinguishes among Australian individuals and explains nearly as much variation as the first component (5.23%). Additional components distinguish among populations nearly as effectively: PC3 (5.12%) shows that the Australian population falls in between UK and NA clusters, and PC4 (4.82%) indicates differences between the NA and other populations (Figure S5).

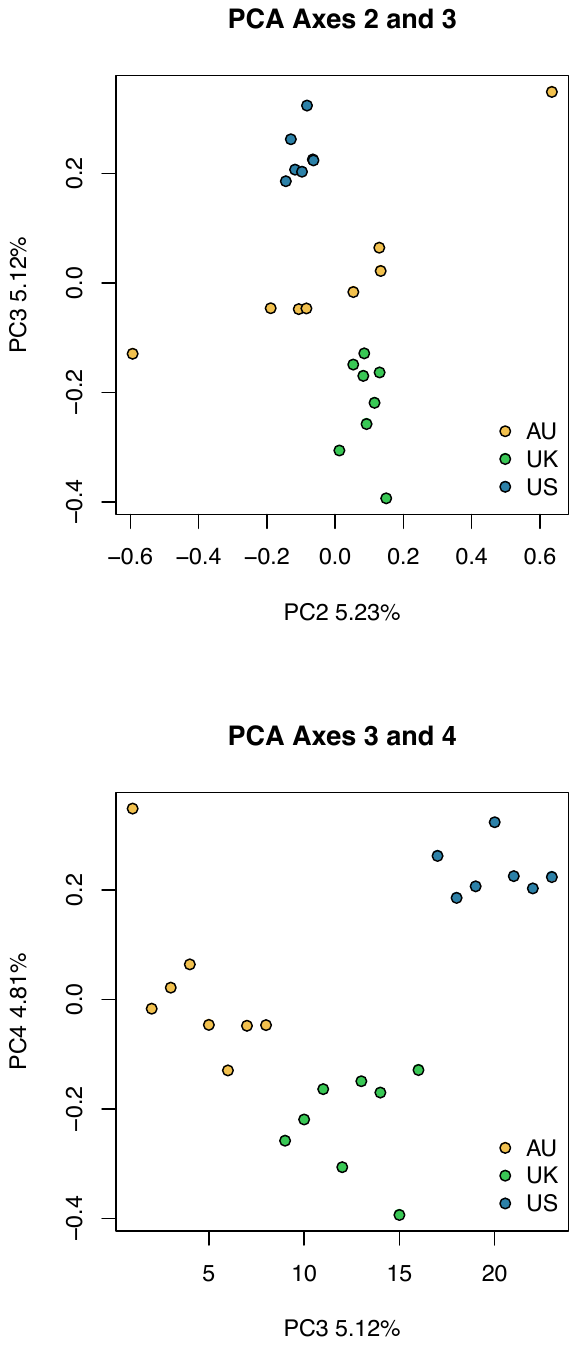

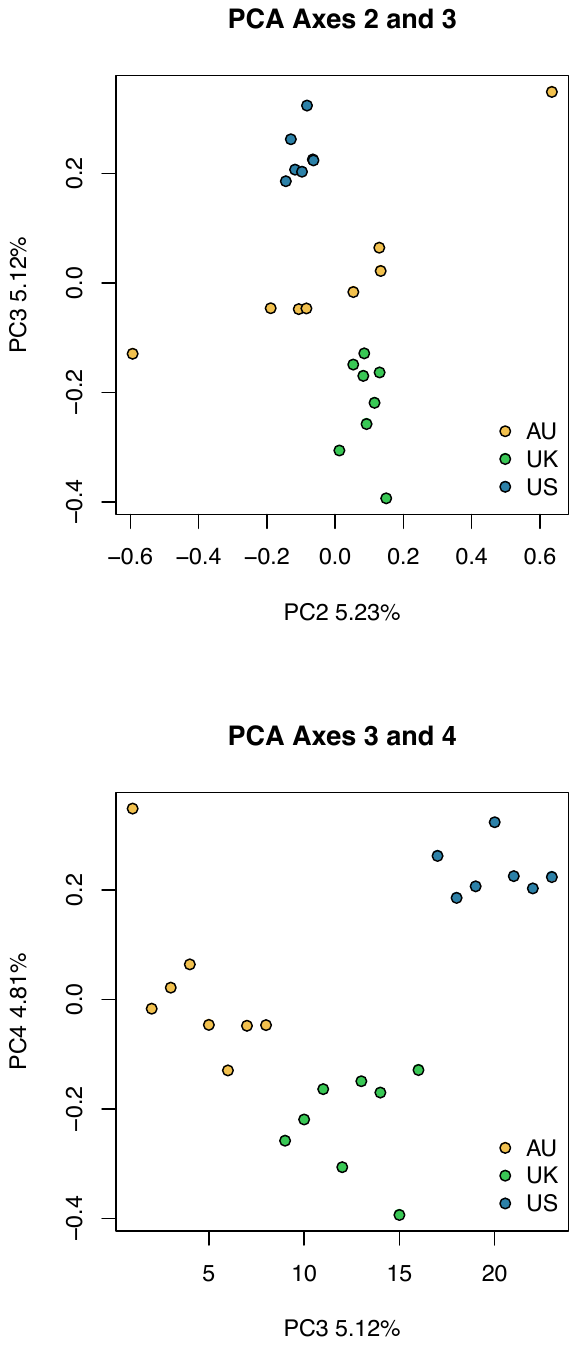

##### **Figure S7.** Additional PCs of a principal component analysis of the variant-called dataset.

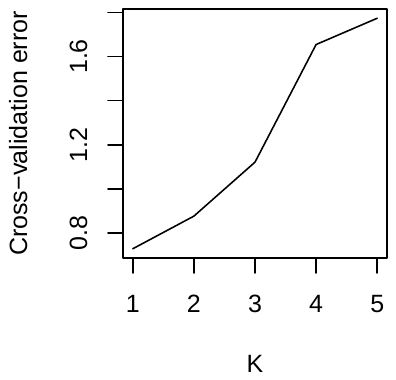

##### **Figure S8.** Cross-validation errors for ADMIXTURE tests of K=1-5.

### 4. Inbreeding and relatedness

##### **Table S2.** Inbreeding statistics (F-statistic) for each individual.

| Individual | F-statistic |
| --- | --- |
| AU1 | 0.32043 |
| AU2 | 0.03098 |
| AU3 | 0.03386 |
| AU4 | 0.05907 |
| AU5 | 0.10412 |
| AU6 | 0.25472 |
| AU7 | 0.06493 |
| AU8 | 0.0591 |
| UK1 | -0.01137 |
| UK2 | -0.02545 |
| UK3 | -0.0242 |
| UK4 | 0.02273 |
| UK5 | -0.02165 |
| UK6 | -0.02398 |
| UK7 | 0.06993 |
| UK8 | -0.01048 |
| US1 | 0.00674 |
| US2 | -0.01957 |
| US3 | 0.48915 |
| US4 | -0.00911 |
| US5 | 0.01501 |
| US6 | -0.00107 |
| US7 | -0.00554 |
| US8 | 0.00771 |

### 5. Genomic architecture of the starling genome

##### **Table S3.** Distance between center of F_ST_ peaks shown in main text and approximate centromere position.

| Chromosome | Centromere position (Mb) | Center of peak (Mb) |
| --- | --- | --- |
| 1 | 98.17 | 105 |
| 1A | 62.54 | 49 |
| 2 | 76.29 | 48 |
| 4 | 16.82 | 23 |
| 4A | 19.79* | 6 |
| 6 | 0.89 | 5.5 |
| Z | 27.51 |  |

*This position is opposite to Volker et al. 2010, so some uncertainty here.

### 6. Gene ontology analyses

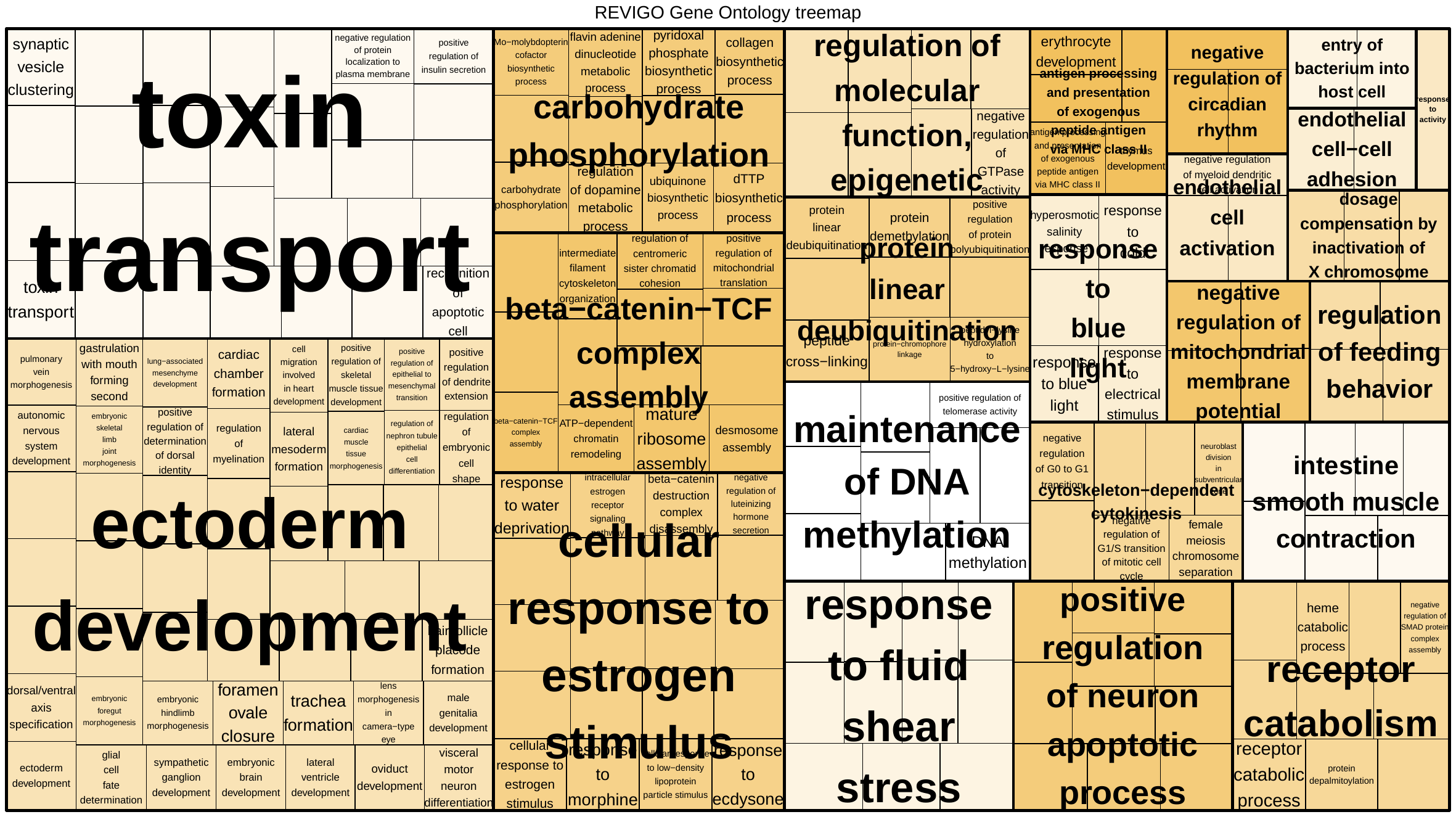

##### **Figure S9.** Common gene ontology (GO) categories represented in Australian vs. UK outliers.

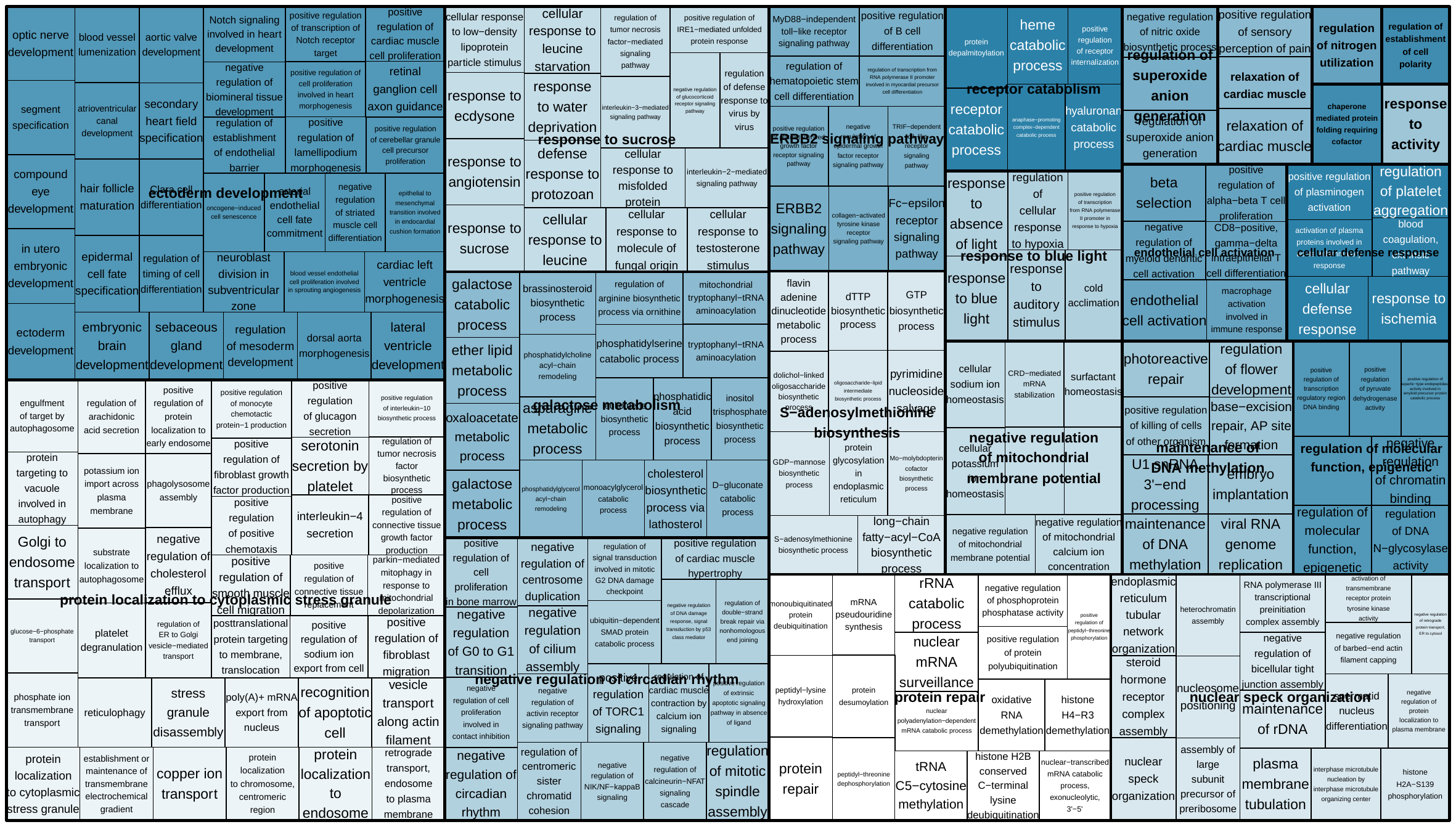

##### **Figure S10.** Common gene ontology (GO) categories represented in NA vs. UK outliers.

### 6. Selection inference at additional F_ST_ peaks

Another region where the evolutionary dynamics at play are unique to one invasion is the peak on Chromosome 1, and although this F_ST_ peak could indicate selection in the NA invasion only, drift is a more parsimonious explanation of these patterns (Figure S11). Where the NA population has differentiated from the UK, Tajima’s *D* oscillates around zero, which suggests that neutral processes drive this differentiation.

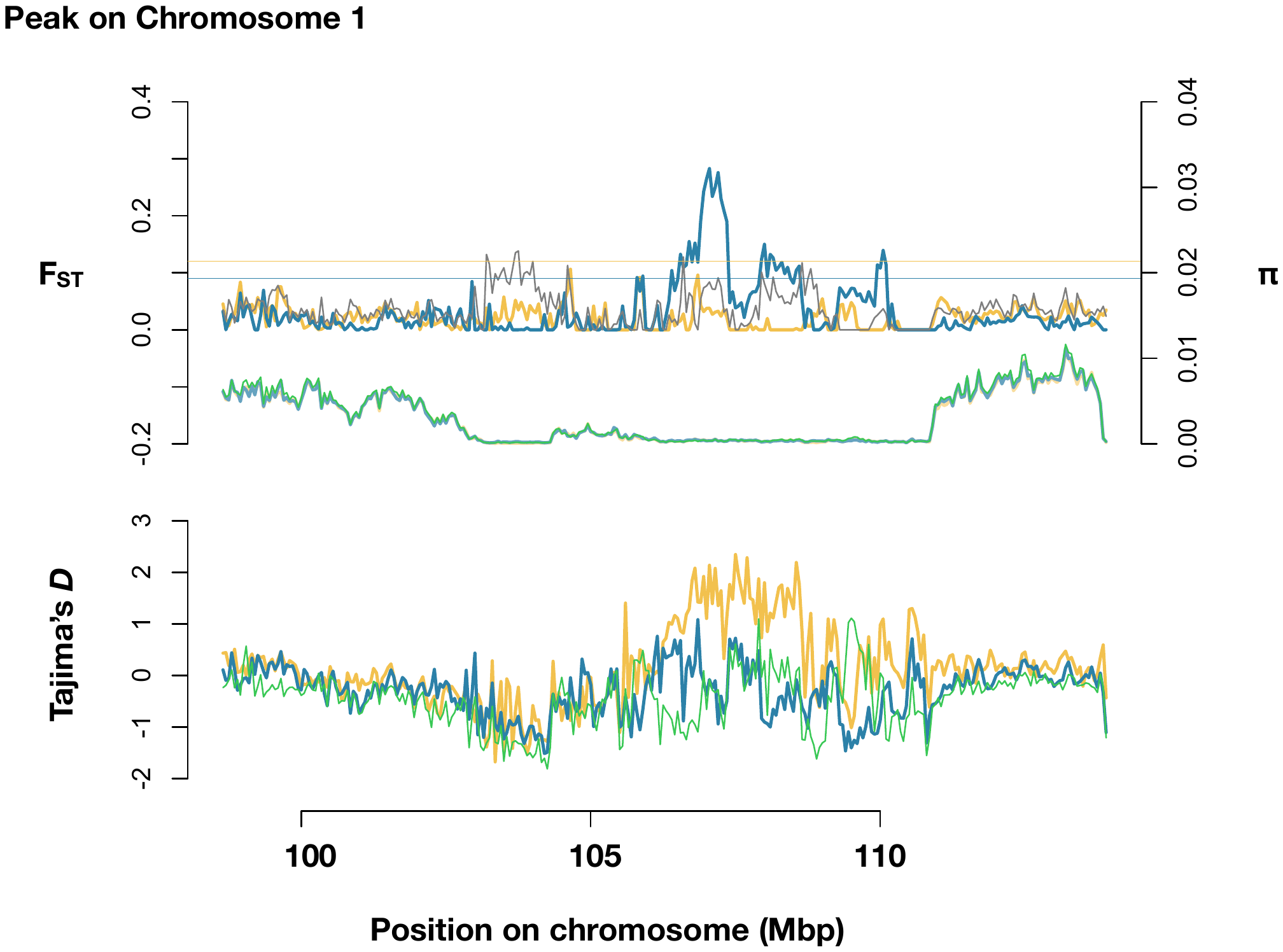

**Figure S11.** Peak on Chromosome 1. 50-kb windowed F_ST_ (top row), π within each population (middle), and Tajima’s *D* within each population (bottom row), centered on the elevated F_ST_ region.

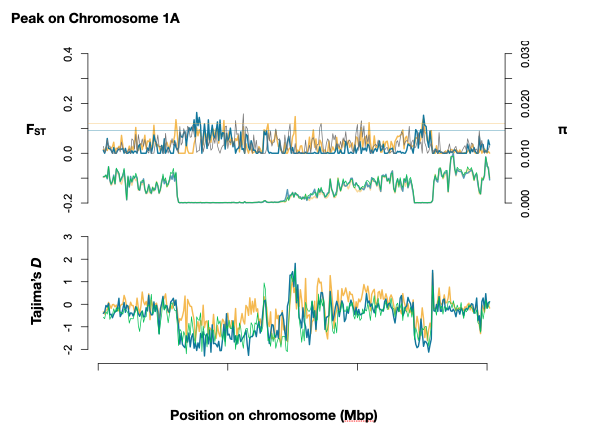

##### **Figure S12.** Peak on Chromosome 1A.

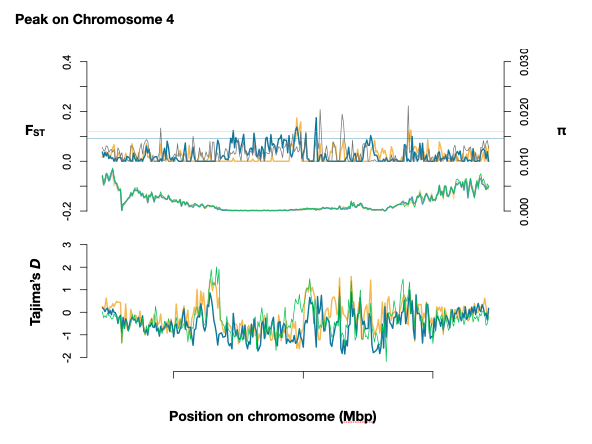

##### **Figure S13.** Peak on Chromosome 4.

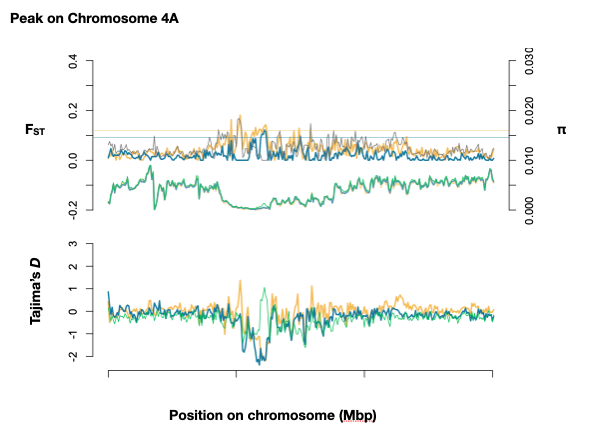

##### **Figure S14.** Peak on Chromosome 4A.
